# DeepCAST-GWAS: Improving the Discovery of Genetic Associations Using Deep Learning-Based Regulatory SNP Prioritization

**DOI:** 10.1101/2025.11.27.690924

**Authors:** Lovro Rabuzin, Konstantin Heep, Sophie Sigfstead, Valentina Boeva

**Author notes:** These authors contributed equally to this work (Lovro Rabuzin, Konstantin Heep).

## Abstract

Genome-wide association studies (GWAS) have uncovered numerous variants linked to complex traits, yet power remains limited by the large multiple testing burden and the inclusion of many variants with minimal regulatory impact. We present Deep learning-based Chromatin Accessibility SNP Targeting for GWAS (DeepCAST-GWAS), a framework that integrates functional annotations derived from deep learning models to improve both the yield and the reliability of GWAS findings. DeepCAST-GWAS uses SNP Activity Difference (SAD) scores from in silico mutagenesis with the Enformer model to estimate the predicted effect of each variant on chromatin accessibility across tissues, allowing statistical testing to focus on variants with stronger regulatory evidence. Using conservative family-wise error rate (FWER) control, DeepCAST-FWER produces fewer associations than existing power-boosting approaches, but the associations it reports replicate in larger cohort GWAS at substantially higher rates. For applications where discovery count is more important, DeepCAST-sFDR increases the number of genome-wide significant findings above baseline GWAS by using the Enformer SAD scores for stratified False Discovery Rate (sFDR) control. DeepCAST-sFDR achieves performance comparable to the strongest competing method, while maintaining reliability on par with a standard GWAS. Subsampling analyses across a wide range of traits confirm these improvements in both sensitivity and replicability. DeepCAST-GWAS offers a principled way to incorporate sequence-based regulatory predictions into population-scale association testing, demonstrating that chromatin accessibility activity scores can improve the stability of GWAS discoveries. The framework is made available at https://github.com/BoevaLab/DeepCAST-GWAS.

## 1 Introduction

Genome-wide association studies have uncovered thousands of loci linked to complex traits, yet most variants have very small effects and only a limited subset surpasses genome-wide significance [1]. With millions of tests and highly polygenic architectures, the Bonferroni standard is often overly conservative, which restricts power, especially for traits with moderate cohort sizes [21]. This has driven interest in external functional annotations that can help prioritize variants with a higher likelihood of regulatory or coding impact. Prior methods have incorporated conservation metrics, chromatin accessibility, histone marks, and composite metascores such as CADD and Eigen through weighted p values, empirical Bayes approaches, or stratified false discovery rate procedures [13, 18, 19, 3, 12, 5, 8]. Although the existing approaches can provide increases in statistical power and locus discovery, large benchmarking efforts show that their improvements tend to be modest, partly because annotation quality and relevance vary across traits and many annotations lack cell-type or tissue specificity [5]. In addition to limited sensitivity, power-boosting methods can introduce instability, with many discoveries failing to replicate in larger cohorts. This has motivated the use of external functional annotations to increase both power and reliability by prioritizing variants with a higher likelihood of regulatory or coding impact [8].

Deep sequence-based regulatory models have raised the resolution at which variant effects can be predicted [27, 11, 15]. In this work, we leverage the Enformer model, which captures the effects of genomic variants on long-range genomic interactions and models cell-type-specific chromatin accessibility, histone marks, and gene expression directly from sequence [2]. Through in silico mutagenesis, Enformer produces SNP Activity Difference (SAD) scores that quantify predicted allelic perturbations across thousands of regulatory outputs. These scores are mechanistic, allele-specific, tissue-resolved, and available genome-wide, making them richer than classical annotations.

We present DeepCAST, a framework that incorporates Enformer-derived regulatory predictions into GWAS while leaving the underlying association test unchanged. DeepCAST has two complementary variants designed for different goals. DeepCAST-FWER applies a pre-GWAS filter that removes variants with negligible predicted regulatory activity in trait-relevant contexts and then performs standard association testing on the remaining variants under a recalibrated family-wise error rate threshold. Because the filter depends only on external annotations, Type I error control is preserved and the resulting associations show substantially improved replication in larger-cohort GWAS. DeepCAST-sFDR instead uses SAD scores as a stratifying variable in a stratified FDR procedure [19]. Coding variants form a separate stratum, and non-coding variants are prioritized based on the magnitude of predicted regulatory activity. This approach increases discovery yield while maintaining reliability similar to baseline GWAS. Both variants enrich the tested hypotheses for functionally supported variants in tissues linked to the phenotype.

These two DeepCAST variants rely on three principles. First, deep learning regulatory models provide more mechanistic, fine-grained predictions than generic annotation databases [2]. Second, pre-GWAS filtering and stratified FDR preserve statistical validity when annotations are external to the association test. Third, limiting the analysis to phenotype-relevant regulatory tracks reduces dilution from irrelevant annotations, a known issue in annotation-weighted methods [5]. Together, these points position DeepCAST as a flexible strategy for improving both the reliability and, when desired, the sensitivity of GWAS by leveraging mechanistic sequence-based regulatory information.

## 2 Methods

### 2.1 Overview

We developed DeepCAST-GWAS, a framework that uses deep learning-derived regulatory predictions to improve discovery sensitivity and replicability of genome-wide association studies. For each variant, the method computes a SAD score from predicted chromatin accessibility changes between alleles across a large set of tissues, and users select the tissues most relevant for the trait. These external predictions can be incorporated in two distinct ways. First, DeepCAST-GWAS can be used conservatively as a filtering approach, maintaining family-wise error rate control (FWER). Here variants with negligible predicted impact in the selected tissues are removed before association testing, while coding variants are always retained (DeepCAST-FWER). Second, DeepCAST-GWAS can be used to increase discovery yield without filtering by assigning each SNP to a stratum based on SAD magnitude and coding status, followed by stratified false discovery control that allocates more permissive thresholds to strata enriched for functional variants (DeepCAST-sFDR). Both modes rely only on external functional predictions and preserve appropriate error control.

### 2.2 Data

We evaluated DeepCAST-GWAS on two complementary sets of GWAS summary statistics. The first dataset consisted of publicly released association results from the Neale Lab UK Biobank project [10]. We restricted our analysis to 636 pathological phenotypes defined by ICD10 codes C02 to R90, which cover a broad range of disease conditions. These GWAS were generated from European ancestry participants using imputed UK Biobank genotypes (HRC, UK10K and 1000 Genomes reference panels) and processed with the Neale Lab quality control pipeline [10], which includes variant-level filtering, removal of related individuals, and standard covariate adjustment. The second dataset consisted of summary statistics used by the KGWAS method [8]. KGWAS provides multiple subsampled GWAS for each of 21 independent traits at a range of sample sizes, with 5 replicates per sample size, in addition to the GWAS conducted on the full cohort (N=374,000). This structure enables systematic replication-based evaluation of method stability across sample sizes.

For functional annotation, we used SAD scores derived from in silico mutagenesis of the Enformer model [2]. SAD scores quantify the predicted change in chromatin accessibility between the alternate and reference allele for each variant across a large collection of tissues. These scores serve as external, association-independent estimates of potential regulatory impact and are used to guide variant filtering and stratification. DeepCAST-GWAS is designed to accept any ISM-derived SNP activity scores as input, not only Enformer-based estimates. Any model that produces allele-specific regulatory predictions through ISM, such as sequence-based CNNs or transformer architectures trained on chromatin accessibility can be substituted to generate the functional annotations required by the method.

### 2.3 Enformer regulatory scores and track selection

For each SNP, we extracted SNP Activity Difference (SAD) values from all available Enformer chromatin accessibility prediction tracks. SAD scores quantify the predicted allele-specific change in accessibility signal across tissues and assays, produced by in silico mutagenesis on the Enformer model [2]. These predictions cover a wide range of cellular contexts, including primary tissues, cell lines and assay types, which allows the method to incorporate biologically relevant regulatory information for diverse traits.

To ensure biological relevance, only tracks corresponding to tissues or cell types implicated in the phenotype should be used. Selecting the appropriate tracks requires expert knowledge of disease biology, developmental origin, and expected regulatory mechanisms. SAD values within each selected track were standardized across SNPs to allow consistent comparison across tracks with heterogeneous scales. For SNP i and track k, the standardized activity was

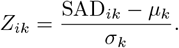

For the large-scale analyses in this study, which involved hundreds of pathological phenotypes and multiple KGWAS subsampling traits, manual annotation of tissue relevance was not feasible. We therefore used an automated procedure in which ChatGPT-4o generated candidate lists of biologically relevant tracks for each phenotype, based on phenotype descriptions and publicly available tissue knowledge. These programmatically generated selections were used for our benchmarking experiments, while we recommend expert curated selections for focused biological studies.

### 2.4 The DeepCAST-FWER framework

DeepCAST-FWER introduces a pre-GWAS filtering stage that restricts association testing to variants with sufficiently strong predicted regulatory impact. For each SNP *i*, we standardize SAD values per track and examine the resulting Z-scores (*Z*_*ik*_). A variant is retained if its maximal absolute standardized activity exceeds a threshold,

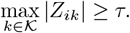

Here *τ* is a user-defined activity threshold. In our experiments we use a fixed value (*τ* = 1), so that only SNPs with standardized activity exceeding this threshold in at least one trait-relevant track are carried forward to association testing. Larger values correspond to more stringent filtering and a smaller set of tested variants, while smaller values allow more permissive inclusion. This threshold can be chosen to reflect the expected regulatory relevance of the trait or the desired balance between sensitivity and specificity.

Let *m*_0_ denote the number of variants in the full GWAS and *m*_*τ*_ the number that remain after DeepCAST filtering. Rather than recomputing a Bonferroni threshold from first principles, we follow common GWAS practice and treat the canonical 5*×*10^*−*8^ significance level as corresponding to the unfiltered hypothesis space. After filtering, we adjust this threshold proportionally to the change in the number of tested variants:

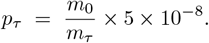

This scaling preserves comparability with standard GWAS analyses while reflecting the reduced multipletesting burden induced by filtering. Because the filtering step depends solely on external regulatory annotations and does not use any association statistics, the adjusted threshold maintains valid family-wise error control. Filtering reduces the effective multiple testing burden and can allow variants with moderate effect sizes to reach the corrected significance threshold, particularly for traits with limited sample sizes or highly polygenic architectures. Since filtering is based entirely on external annotations, the subsequent Bonferroni correction maintains valid FWER control.

DeepCAST-FWER is compatible with any downstream GWAS method. It can be used with linear or logistic regression, linear mixed model association tests, or widely used software such as BOLT-LMM, SAIGE, Regenie, or fastGWA. The framework provides a simple and computationally efficient way to incorporate functional priors into stringent genome-wide significance testing while preserving strict error control.

### 2.5 The DeepCAST-sFDR framework

Many complex traits show different regulatory architectures for coding and non-coding variation, and these differences can be leveraged to improve association power. Stratified false discovery rate control (sFDR) provides a principled way to account for such heterogeneity by grouping hypotheses into strata that are expected to differ in their proportion of non-null effects, and then applying FDR control within each group [19]. DeepCAST-sFDR implements this strategy by stratifying variants according to their predicted functional relevance and performing the Benjamini–Hochberg (BH) FDR control separately within each stratum.

To construct strata, DeepCAST-sFDR uses two sources of prior information:

1. **Coding stratum**. All variants annotated as exonic, reflecting their higher prior probability of functional impact [4].
2. **Non-coding strata based on regulatory activity**. For each non-coding variant, we compute the mean standardized SNP Activity Difference (SAD) across the phenotype-specific set of Enformer chromatin accessibility tracks. Variants are then assigned to non-overlapping SAD activity bins using fixed thresholds

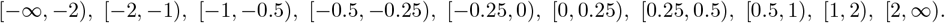

These bins reflect increasing predicted regulatory activity and therefore differing prior probabilities of association.

Once strata are defined, DeepCAST-sFDR follows the “fixed FDR” stratified procedure of Sun et al. [19]. Specifically, within each stratum we apply the BH procedure at the common FDR level *q*. This yields stratum-specific adjusted *p* values, and variants with adjusted *p < q* in their respective stratum are declared significant. Because each stratum receives the full nominal FDR level, strata believed to contain a greater proportion of true associations (for example, coding variants or non-coding variants with high SAD activity) tend to yield more discoveries, while strata with weaker regulatory support contribute fewer.

This stratified BH approach preserves control of the FDR within each stratum and aligns with the theoretical guarantees described by Sun et al. under the fixed-FDR paradigm, while enabling DeepCAST-sFDR to prioritize variants by predicted regulatory relevance without removing any variants from analysis. As a result, DeepCAST-sFDR provides a complementary alternative to DeepCAST-FWER, redistributing statistical power across biologically informed strata and improving sensitivity in high-priority regulatory regions.

## 3 Results

### 3.1 DeepCAST-FWER Improves Discovery Power Over Findor

To assess the ability of DeepCAST-FWER to increase statistical power in genome-wide association studies, we benchmarked the method against FINDOR, a widely used functional-annotation-based approach designed to improve GWAS power through variant prioritization [12], across a broad set of complex traits. We first compared the number of distinct loci recovered by each approach relative to the baseline GWAS. Across traits with comparable baseline signal levels, DeepCAST-FWER identified more genome-wide significant loci than FINDOR (Fig. 2A). Across 336 phenotypes, DeepCAST-FWER identified more genome-wide significant loci than FINDOR in 164 phenotypes (49 percent). These gains were particularly pronounced for phenotypes with modest numbers of baseline associations. When restricting the analysis to traits with fewer than fifteen baseline loci, DeepCAST-FWER showed larger increases in locus discovery, with a mean increase of 0.3 loci, indicating that it benefits underpowered phenotypes where FINDOR provides limited improvements (Fig. 2B).

**Fig. 1.**
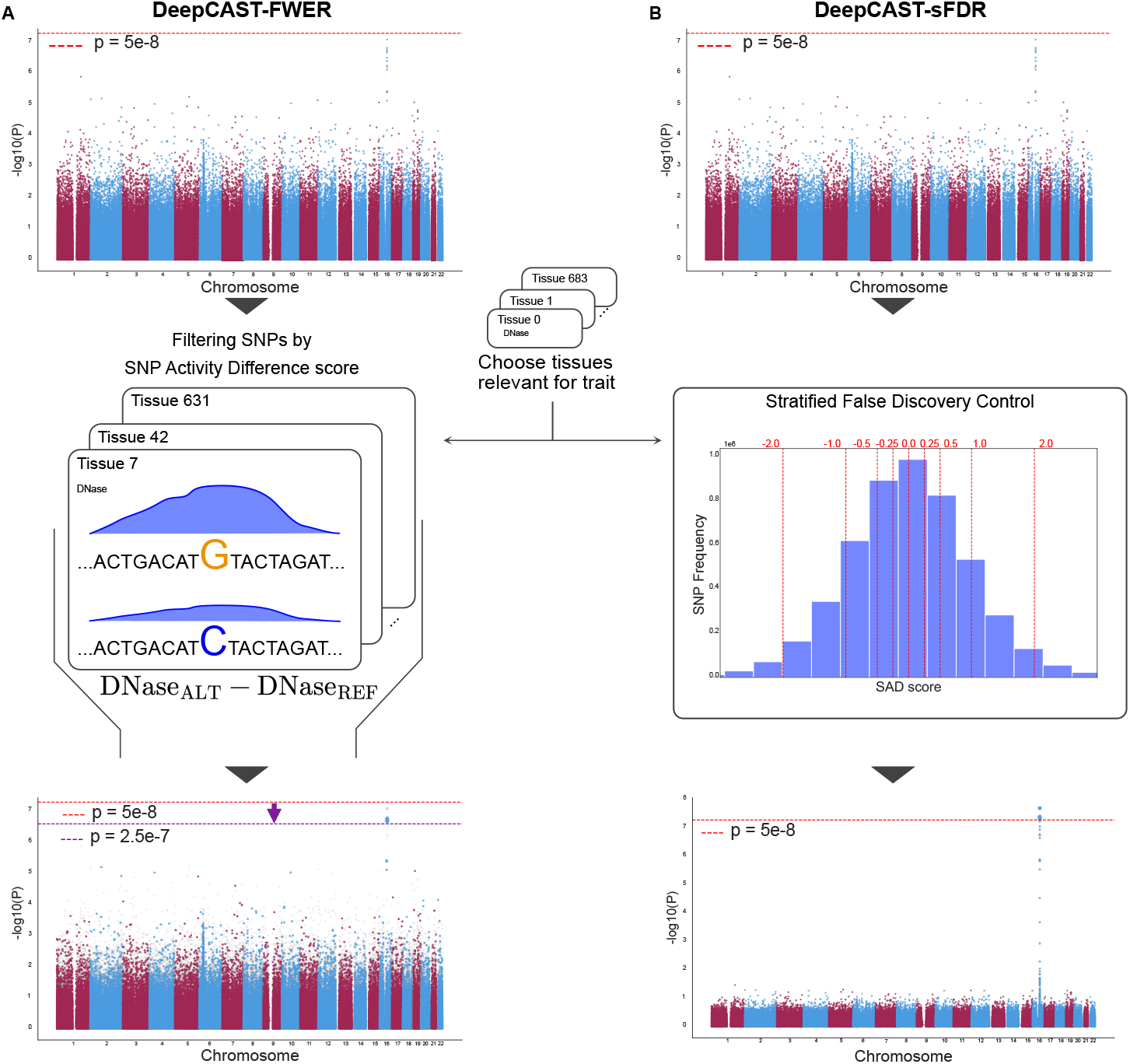
Overview of the DeepCAST-GWAS workflow. **A**, DeepCAST-FWER. A standard GWAS approach tests all variants genome-wide, which leads to a large multiple testing burden. DeepCAST-GWAS introduces an external, prediction-based filtering step that does not use any information from the association statistics. For each SNP, the model uses a SNP Activity Difference (SAD) score based on the predicted change in chromatin accessibility between the alternate and reference alleles across multiple tissues. The user selects tissues relevant for the trait, and non-coding SNPs with negligible predicted impact across these tissues are removed before association testing, improving the reliability of the resulting associations. This filtering relies only on sequence-based regulatory predictions and is entirely independent of the GWAS *p*-values. **B**, DeepCAST-sFDR. Remaining SNPs are grouped into strata defined by their SAD score magnitude and by coding status. Stratified false discovery control is then applied to increase discovery yield while maintaining error control. This yields a final association scan that concentrates power on SNP groups with higher predicted regulatory impact while maintaining proper genome-wide control of false discoveries.

**Fig. 2.**
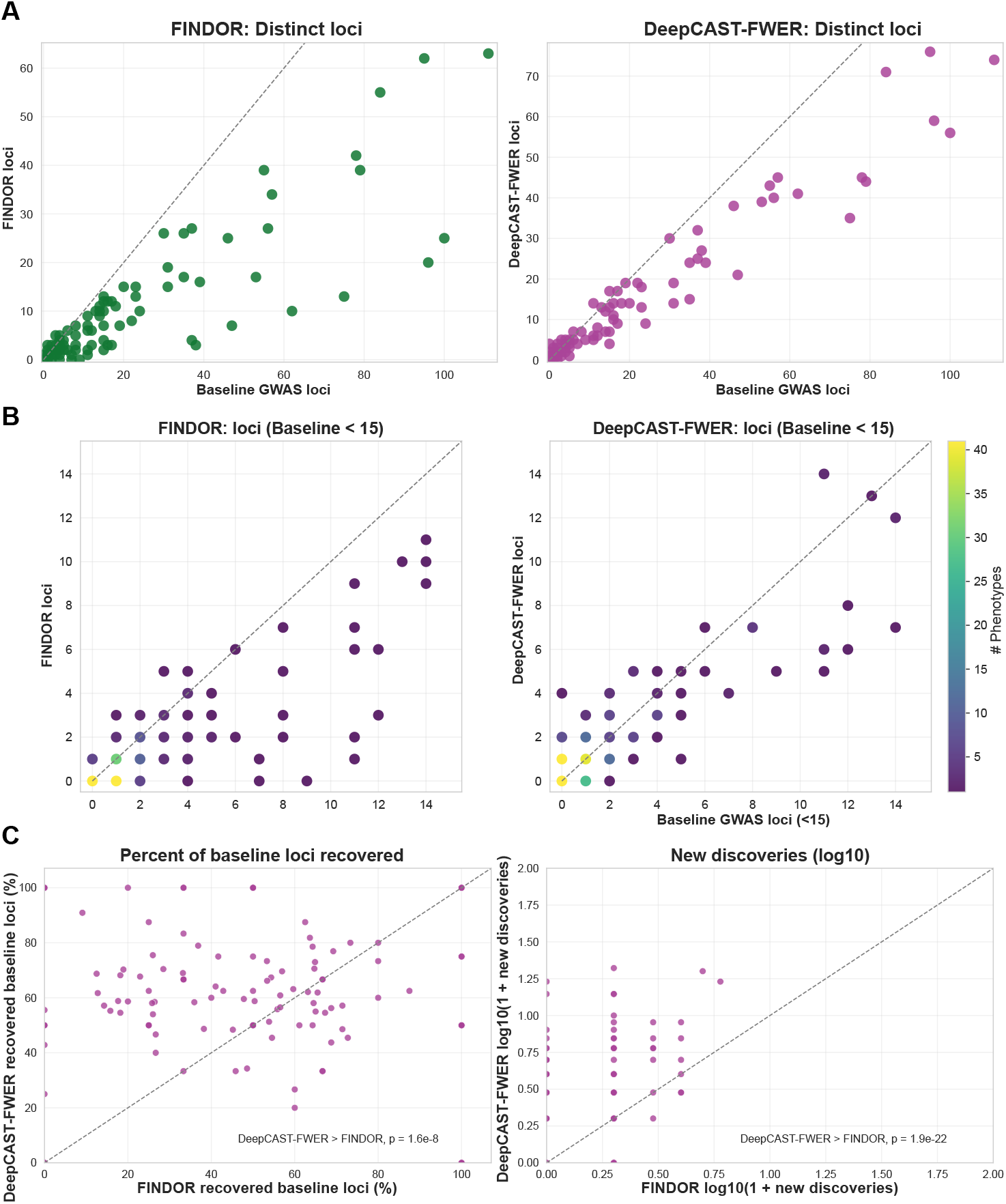
Comparison of DeepCAST-FWER with FINDOR across a broad set of complex traits. **A**, Scatter plots of distinct loci discovered by each method relative to the baseline GWAS show that DeepCAST-FWER consistently identifies more loci than FINDOR for phenotypes with similar baseline signal levels. Each dot corresponds to a GWAS study for a specific phenotype. **B**, For phenotypes with fewer than fifteen baseline loci, DeepCAST-FWER yields larger increases in locus discovery than FINDOR, with points colored by the number of phenotypes contributing to each count. **C**, Direct method-to-method comparisons reveal that DeepCAST-FWER recovers a higher fraction of baseline loci (*p* = 1.6e-8) and produces substantially more new discoveries (*p* = 1.9e-22) than FINDOR.

We next performed direct method-to-method comparisons to quantify both recovery of known loci and identification of new associations. DeepCAST-FWER recovered a significantly higher fraction of baseline GWAS loci than FINDOR (*p* = 1.6e-08, one-sided Wilcoxon signed-rank test), demonstrating that its discoveries are more likely to align with established genetic signals. The method also produced substantially more new loci at genome-wide significance (*p* = 1.9e-22, one-sided Wilcoxon signed-rank test), highlighting its ability to uncover additional trait-associated regions that FINDOR fails to detect (Fig. 2C).

Together, these results show that DeepCAST-FWER provides a robust power increase over FINDOR while maintaining strong concordance with baseline GWAS signals, particularly in settings where sample sizes or inherent effect sizes limit traditional discovery power.

### 3.2 DeepCAST-sFDR Improves Sensitivity Under FDR Control

Next, we evaluated DeepCAST-sFDR relative to an annotation-based stratified FDR procedure using CADD scores [13, 17] as the stratifying variable. Following the strategy of Gao et al. [5], we formed two strata: the top 5% of variants ranked by the CADD score and the remaining 95%, and applied FDR control separately within each stratum (CADD-sFDR). This comparison tests whether chromatin accessibility predictions alone provide a more informative basis for prioritization than widely used composite annotations.

Across a wide range of complex traits, DeepCAST-sFDR identified more distinct loci than CADD-sFDR when benchmarked against the baseline GWAS (Fig. 3A). DeepCAST-sFDR identified more loci than CADD-sFDR in 9.8 percent of phenotypes. These improvements were not restricted to highly powered traits. Among phenotypes with fewer than fifteen baseline loci, DeepCAST-sFDR showed higher gains in locus discovery, indicating a particular advantage in settings where the available association signal is limited (Fig. 3B).

**Fig. 3.**
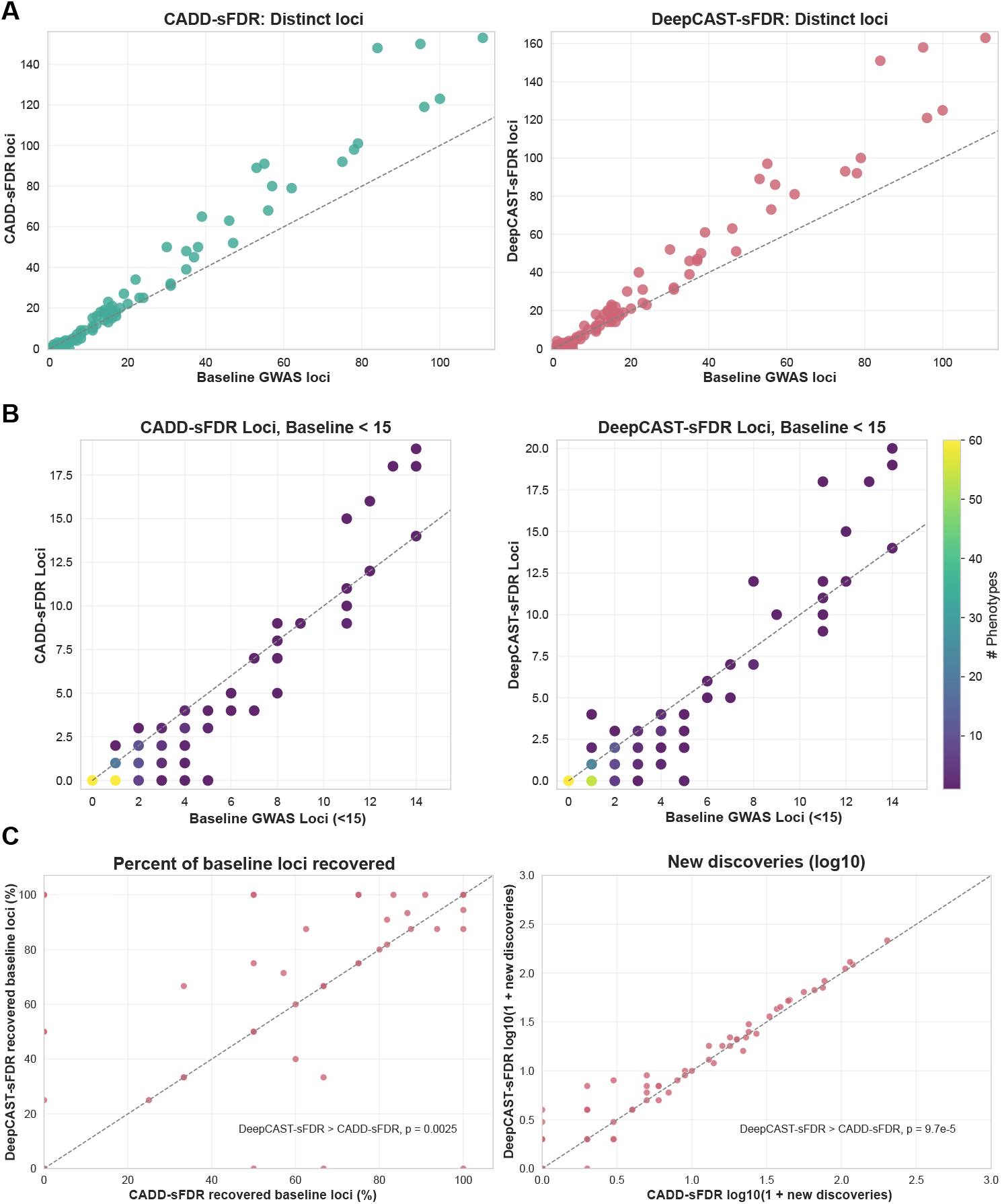
Comparison of DeepCAST-sFDR with CADD-sFDR across diverse complex traits. **A**, Scatter plots of distinct loci discovered by each method relative to the baseline GWAS show that DeepCAST-sFDR yields larger gains than CADD-sFDR. **B**, Among traits with fewer than fifteen baseline loci, DeepCAST-sFDR identifies more loci than CADD-sFDR, with points colored by the number of phenotypes contributing to each bin. **C**, Direct method-to-method comparisons indicate that DeepCAST-sFDR recovers a greater share of baseline loci (*p* = 0.0025) and produces more new discoveries (*p* = 9.7e-5) than CADD-sFDR, highlighting its improved power in the FDR setting.

To compare the two stratification strategies more directly, we examined replication of baseline GWAS loci and the discovery of additional associations. DeepCAST-sFDR recovered a significantly greater fraction of the baseline loci than CADD-based stratification (*p* = 0.0025, one-sided Wilcoxon signed-rank test), demonstrating improved alignment with established genetic signals. It also identified more new loci at the same FDR level (*p* = 9.7e-5, one-sided Wilcoxon signed-rank test), confirming that chromatin accessibility–based stratification provides higher sensitivity than CADD-based stratification under FDR control (Fig. 3C).

Together, these results show that DeepCAST-sFDR offers stronger gains in both recovery and expansion of trait-associated loci than a stratified FDR procedure based on CADD scores, highlighting the advantage of using sequence-derived, tissue-resolved regulatory predictions for variant prioritization.

### 3.3 Replication-Focused and Discovery-Focused Performance in Subsampled Cohorts

To evaluate not only discovery but also reliability, we conducted a replication-based subsampling analysis across five cohort sizes ranging from one thousand to ten thousand individuals. We compared DeepCAST-FWER and DeepCAST-sFDR with GWAS, FINDOR, KGWAS, and CADD-sFDR using the full cohort GWAS as the gold standard. DeepCAST-FWER consistently produced associations with the highest proportion of signals that replicated in the full cohort across most sample sizes (Fig. 4A). Although DeepCAST-FWER yielded fewer total discoveries than power boosting methods such as FINDOR and KGWAS, its findings were more reliable. DeepCAST-FWER consistently achieved the highest proportion of replicated findings across most sample sizes, demonstrating robustness in underpowered settings. In contrast, DeepCAST-sFDR provided a complementary balance between sensitivity and precision. It increased the number of replicated associations beyond the baseline GWAS and performed similarly to KGWAS in terms of replicated discovery count, while maintaining replication in the full cohort that closely matched both GWAS and KGWAS.

**Fig. 4.**
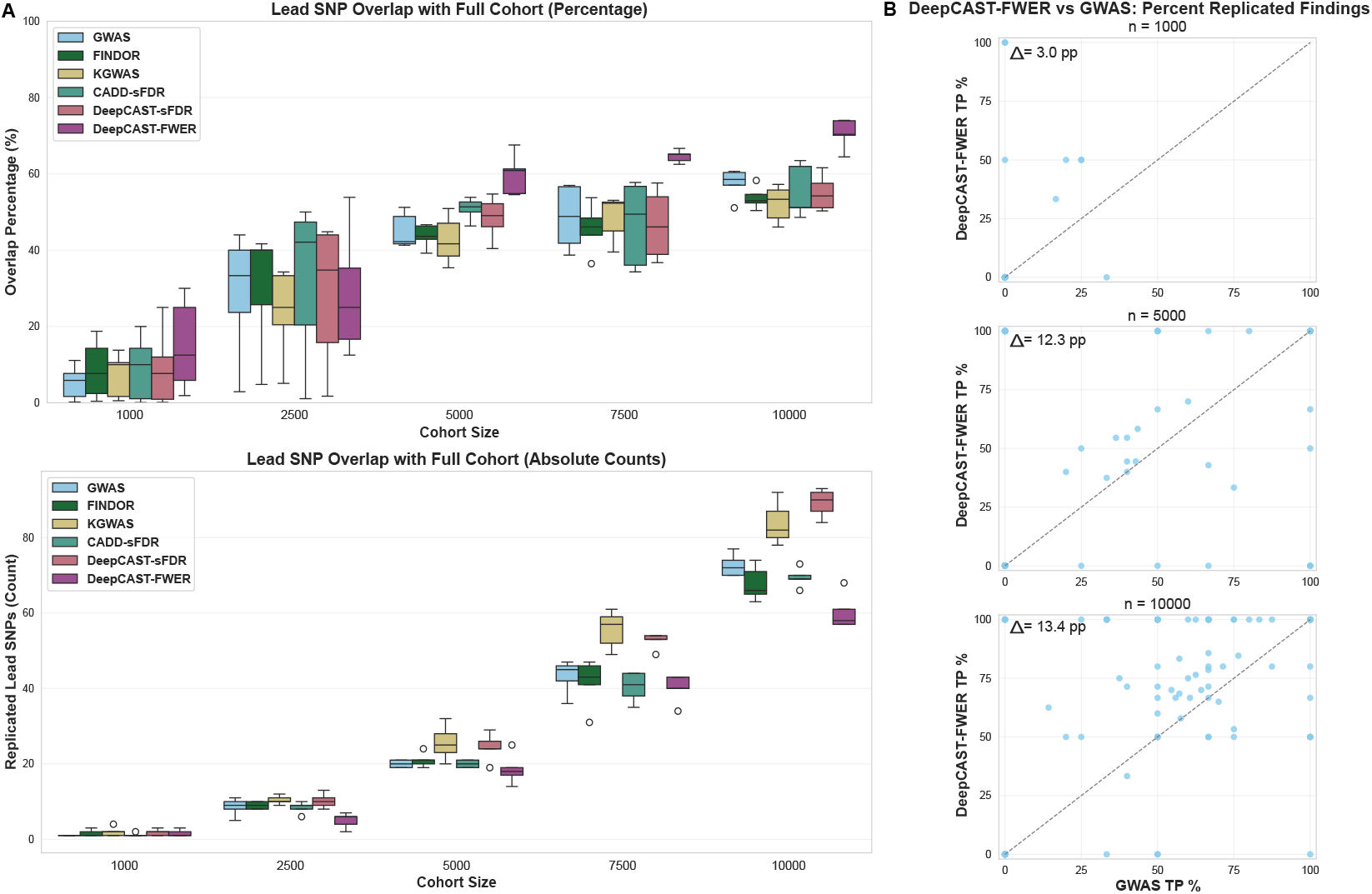
Subsampling analysis comparing DeepCAST-FWER with GWAS, FINDOR, KGWAS, CADD- sFDR, and DeepCAST-sFDR. **A**, Across cohort sizes, DeepCAST-FWER produces associations with a higher proportion of signals that replicate in the full cohort GWAS, although it yields fewer total discoveries than power boosting methods such as FINDOR and KGWAS. DeepCAST-sFDR provides an alternative option for users prioritizing discovery count, offering more associations than the baseline GWAS and achieving performance comparable to KGWAS while retaining similar reliability. **B**, Pairwise subsampling comparisons across phenotypes show variable performance on individual traits, yet DeepCAST-FWER exhibits a clear overall advantage, achieving higher replicated true positive rates than GWAS when aggregated across sample sizes.

The pairwise subsampling comparisons further illustrate how the methods behave across diverse traits and replicate subsets. Although performance varied between individual phenotypes, DeepCAST-FWER showed a consistent overall advantage, producing a higher proportion of replicated true positives than the baseline GWAS when the results were aggregated across all sample sizes and subsampling replicates (Fig. 4B).

### 3.4 DeepCAST-GWAS Identifies Novel Loci and Prioritizes Functionally Supported Variants

To illustrate how DeepCAST-GWAS integrates chromatin accessibility information to refine association signals, we examined specific loci identified in representative traits.

For atrioventricular and left bundle-branch block, DeepCAST-FWER discovered a genome-wide significant locus on chromosome 3 that the baseline GWAS did not detect (Fig. 5A). The lead variant, rs6781009, lies within a VISTA-annotated cardiac enhancer [14, 23] and overlaps DNase and H3K27ac ChIP-seq peaks from left-ventricle tissue [16, 26] and is positioned between the genes *EXOG* and *SCN5A. EXOG* encodes a mitochondrial 5’-3’ exonuclease that participates in the repair and maintenance of mitochondrial DNA [20], and has been previously associated with electrocardiography scores [7]. Although EXOG has not been directly tied to cardiac conduction disorders, the mitochondrial stress and apoptosis pathways it participates in may influence conduction tissue health and vulnerability. In contrast, SCN5A encodes a major cardiac sodium channel, which is directly responsible for the inward sodium current that initiates the cardiac action potential upstroke and enables impulse propagation through the atrioventricular node [22]. Mutations or dysregulation of SCN5A have been firmly implicated in conduction block, arrhythmias, and bundle-branch block phenotypes [22]. The proximity of the variant to these genes, particularly SCN5A, strengthens the plausibility that this locus modifies cardiac conduction via regulatory effects on one or both of these genes. In a second example, we applied DeepCAST-FWER to lung cancer summary statistics. Both baseline GWAS and DeepCAST identified the same broad susceptibility region, but the methods prioritized different lead variants. DeepCAST selected rs72738786 in the 3’-UTR of *HYKK*, whereas the baseline reported rs11852372 lies in an intronic region of the same gene (Fig. 5B). Variants in 3’-UTRs commonly influence gene regulation through effects on mRNA stability, microRNA binding or transcript processing [6, 25]. Since the baseline reported variant does not overlap known regulatory marks or splicing motifs, the positional context of the DeepCAST-prioritized variant provides a more plausible functional mechanism. This highlights the method’s ability to refine association signals even when overall locus discovery overlaps with standard GWAS.

**Fig. 5.**
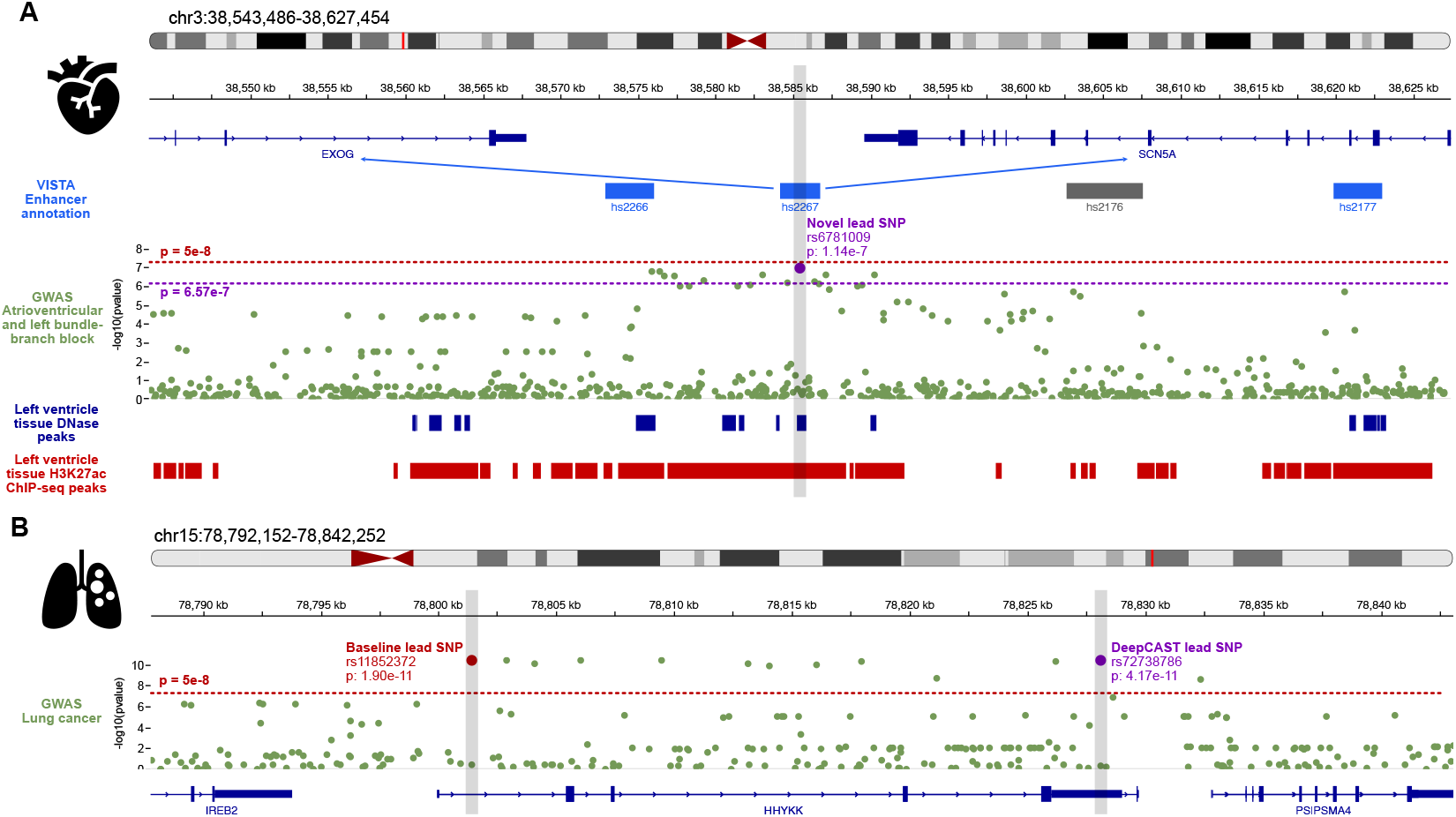
DeepCAST-GWAS discovers novel loci and prioritizes variants more likely to be functional. **A**, Application of DeepCAST-FWER to atrioventricular and left bundle-branch block GWAS highlights a novel locus on chromosome 3 that the baseline GWAS does not detect. DeepCAST identifies rs6781009 within an enhancer active in cardiac tissue and overlapping left ventricle DNase and H3K27ac ChIP-seq peaks, positioned between the genes *EXOG* and *SCN5A*. **B**, Application to lung cancer GWAS recovers the same susceptibility region as the baseline scan but selects a different lead variant: DeepCAST prioritizes rs72738786 in the 3’-UTR of *HYKK*, while the baseline identifies rs11852372 located in an intron of *HYKK*. Together, these examples illustrate how DeepCAST can reveal novel signals and refine lead variant selection by integrating regulatory annotations.

These case studies demonstrate that DeepCAST-GWAS can reveal novel loci associated with traits and improve lead variant prioritization by incorporating sequence-based regulatory predictions, thereby enhancing the biological relevance of the GWAS findings.

## 4 Discussion

DeepCAST provides a flexible framework for improving the reliability and interpretability of genome-wide association studies by integrating deep learning derived regulatory predictions into standard association workflows. DeepCAST-FWER uses mechanistic annotations to filter variants prior to testing, reducing the multiple testing burden while preserving strict family-wise error control. In practice, DeepCAST-FWER yields fewer total associations than power-boosting methods such as FINDOR or KGWAS, but the associations it identifies replicate at substantially higher rates in full-cohort analyses. This makes DeepCAST-FWER well suited for applications where the primary objective is to identify a compact and highly reliable set of candidate loci, rather than to maximize the number of discoveries.

DeepCAST-sFDR complements this conservative mode with a softer prioritization strategy. Instead of filtering variants, it stratifies the hypothesis space by coding status and predicted regulatory activity, and applies false discovery rate control independently within each stratum. This approach increases the total number of discoveries relative to baseline GWAS and performs comparably to KGWAS, while maintaining reliability that is similar to or better than unstratified FDR control. DeepCAST-sFDR therefore offers an appealing option for settings where a larger set of associations is desirable but robustness must be preserved. A key strength of DeepCAST lies in its use of tissue-resolved regulatory predictions that model the biochemical consequences of sequence variation. This distinguishes DeepCAST from classical annotation-weighted approaches that rely on generic or aggregated annotations, which often lack trait specificity and have shown modest or inconsistent gains when integrated into GWAS [5]. By selecting regulatory tracks relevant to each phenotype, DeepCAST focuses statistical power on the biological contexts most likely to mediate genetic effects, which improves locus prioritization and enhances the interpretability of significant associations.

Although newer sequence-to-function models such as Borzoi [15] are now available, we used Enformer because, to the best of our knowledge, it remains among the strongest publicly accessible predictors of DNase and chromatin accessibility, the regulatory modality directly used to prioritize variants in our approach. This choice ensures that DeepCAST leverages high-quality accessibility predictions, which are central to our filtering and stratification procedures. The selection of regulatory tracks and tissues is another important factor in DeepCAST’s performance. While our automated procedure for large-scale benchmarking performed well across hundreds of phenotypes, different track selections may shift sensitivity for specific traits. Developing principled, data-driven strategies to infer phenotype-relevant tissues represents a natural direction for future refinement.

Several additional extensions could further enhance DeepCAST. Incorporating additional modalities predicted by deep learning models, such as histone modifications, promoter–enhancer interactions, or RNA abundance, may improve performance for traits driven by diverse regulatory mechanisms. Learning adaptive, data-driven strata for sFDR or tuning filtering thresholds for FWER control could refine the balance between sensitivity and reliability. Furthermore, integrating DeepCAST with fine-mapping [28], burden testing [9], or polygenic prioritization frameworks [24] may broaden its utility across varied study designs.

Overall, DeepCAST offers a simple, modular, and statistically principled strategy for improving both the stability and biological relevance of GWAS findings. By combining classical statistical guarantees with mechanistic, allele-specific regulatory predictions from modern deep learning models, DeepCAST provides a practical approach for prioritizing genetic variants in trait-relevant regulatory contexts and extracting more informative biological insights from association studies.

## 5 Code availability

DeepCAST-GWAS is implemented in Python and is available as an open-source repository at https://github.com/BoevaLab/DeepCAST-GWAS. The repository includes a comprehensive README with full usage instructions, as well as example SLURM scripts for running the pipeline in high-performance computing (HPC) environments. The code covers the complete workflow, including data preprocessing, integration of Enformer-derived SAD scores, variant filtering, stratified FDR analysis, and subsamplingbased replication benchmarking.

