## Appendix for "DeepCAST-GWAS: Improving the Discovery of Genetic Associations Using Deep Learning-Based Regulatory SNP Prioritization"

### Appendix A Supplementary figures

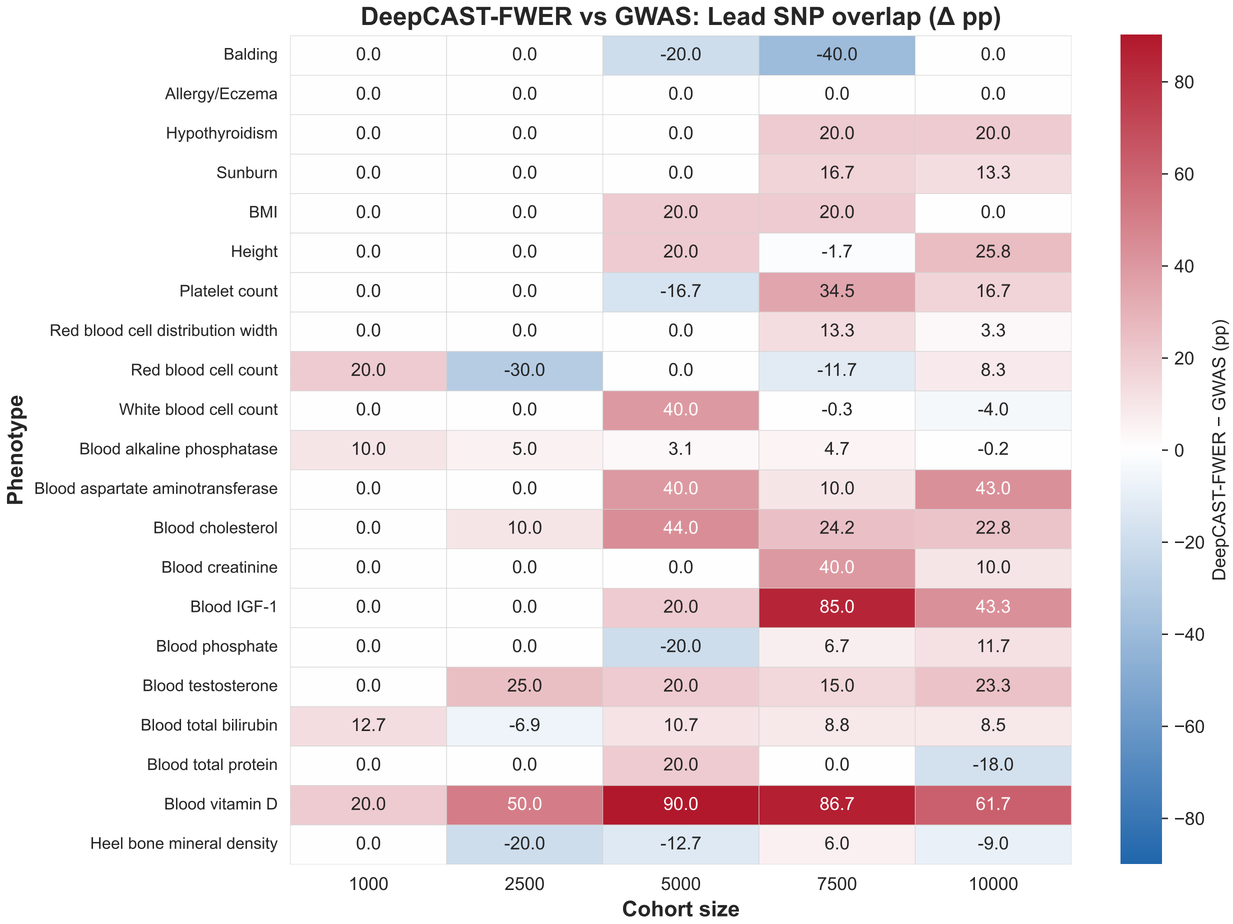


| Phenotype | 1000 | 2500 | 5000 | 7500 | 10000 |
|---|---|---|---|---|---|
| Balding | 0.0 | 0.0 | -20.0 | -40.0 | 0.0 |
| Allergy/Eczema | 0.0 | 0.0 | 0.0 | 0.0 | 0.0 |
| Hypothyroidism | 0.0 | 0.0 | 0.0 | 20.0 | 20.0 |
| Sunburn | 0.0 | 0.0 | 0.0 | 16.7 | 13.3 |
| BMI | 0.0 | 0.0 | 20.0 | 20.0 | 0.0 |
| Height | 0.0 | 0.0 | 20.0 | -1.7 | 25.8 |
| Platelet count | 0.0 | 0.0 | -16.7 | 34.5 | 16.7 |
| Red blood cell distribution width | 0.0 | 0.0 | 0.0 | 13.3 | 3.3 |
| Red blood cell count | 20.0 | -30.0 | 0.0 | -11.7 | 8.3 |
| White blood cell count | 0.0 | 0.0 | 40.0 | -0.3 | -4.0 |
| Blood alkaline phosphatase | 10.0 | 5.0 | 3.1 | 4.7 | -0.2 |
| Blood aspartate aminotransferase | 0.0 | 0.0 | 40.0 | 10.0 | 43.0 |
| Blood cholesterol | 0.0 | 10.0 | 44.0 | 24.2 | 22.8 |
| Blood creatinine | 0.0 | 0.0 | 0.0 | 40.0 | 10.0 |
| Blood IGF-1 | 0.0 | 0.0 | 20.0 | 85.0 | 43.3 |
| Blood phosphate | 0.0 | 0.0 | -20.0 | 6.7 | 11.7 |
| Blood testosterone | 0.0 | 25.0 | 20.0 | 15.0 | 23.3 |
| Blood total bilirubin | 12.7 | -6.9 | 10.7 | 8.8 | 8.5 |
| Blood total protein | 0.0 | 0.0 | 20.0 | 0.0 | -18.0 |
| Blood vitamin D | 20.0 | 50.0 | 90.0 | 86.7 | 61.7 |
| Heel bone mineral density | 0.0 | -20.0 | -12.7 | 6.0 | -9.0 |

**Fig. 1. DeepCAST-FWER increases replication of lead SNPs in the full-cohort GWAS compared to standard GWAS.** The heatmap displays the difference in the proportion of lead SNPs replicated in the full cohort ($\Delta$ percentage points). Positive values (red) denote improved overlap under DeepCAST-FWER, while negative values (blue) indicate reduced overlap. DeepCAST-FWER improves replication across many traits, especially for vitamin D, and IGF1.

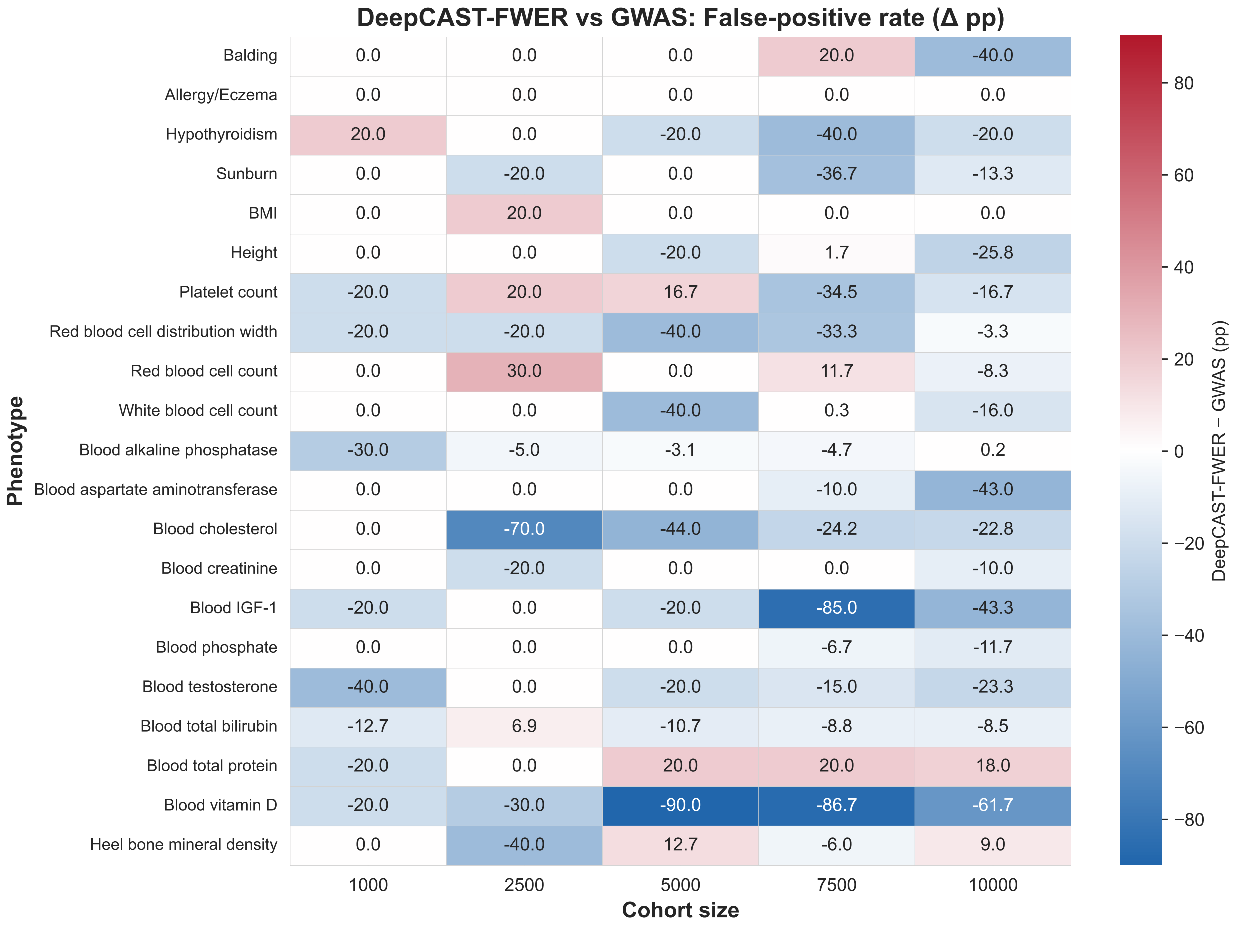


**Fig. 2. DeepCAST-FWER reduces false positive rates relative to standard GWAS across subsampled cohort sizes.** For each phenotype and cohort size, the heatmap shows the change in the percentage of lead SNPs that fail to replicate in the full-cohort GWAS ($\Delta$ percentage points). Negative values (blue) indicate fewer false positives under DeepCAST-FWER, while positive values (red) indicate more. DeepCAST-FWER frequently lowers false positive rates, particularly for biochemical traits and medium-sized cohorts.

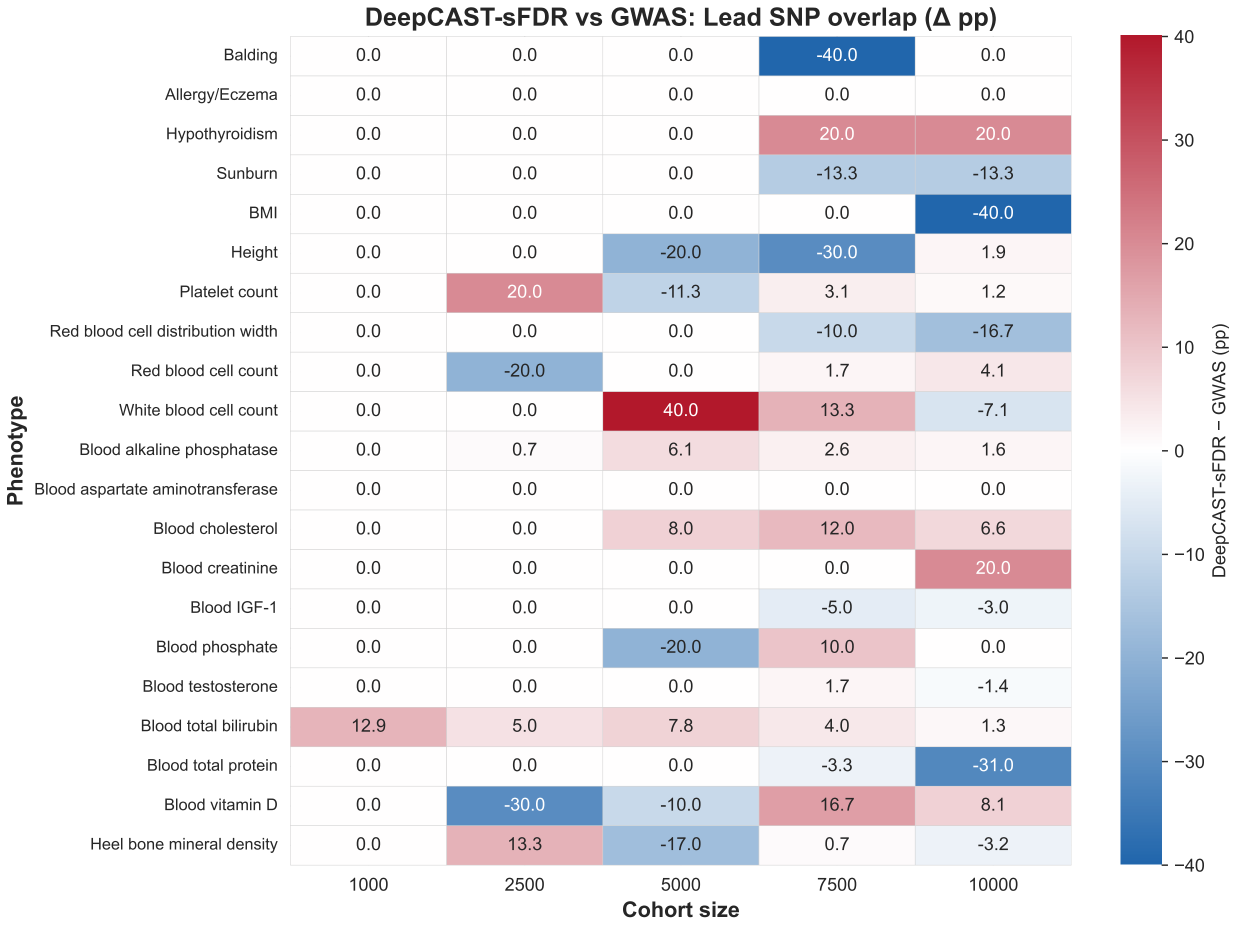


**Fig. 3. DeepCAST-sFDR achieves replication rates comparable to GWAS.** This heatmap reports the change in replicated lead SNPs ($\Delta$ percentage points) relative to the full-cohort GWAS. Improvements (red) occur in several phenotypes and sample sizes, including total bilirubin, vitamin D, and selected cardiometabolic traits, while decreases (blue) appear in others.

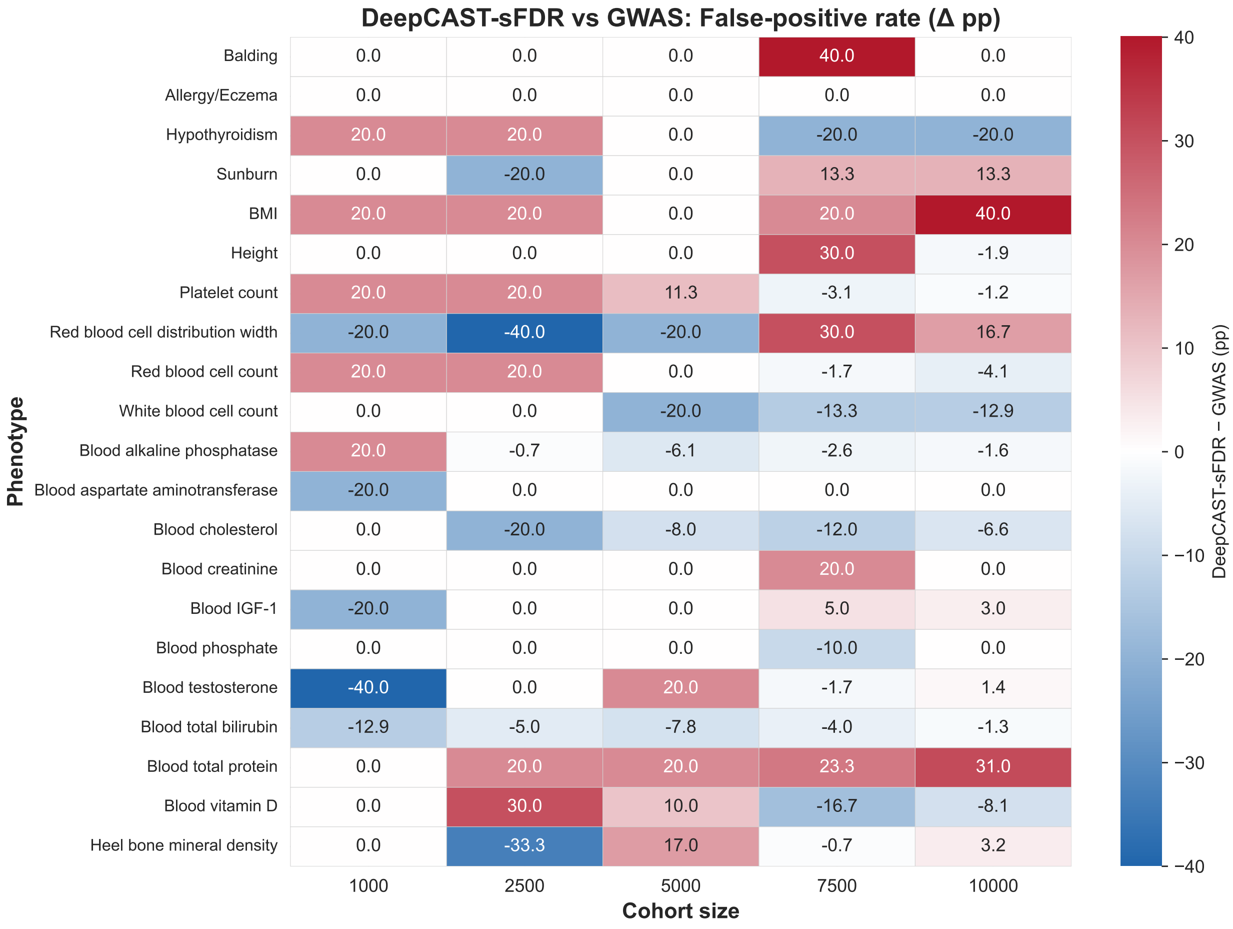


**Fig. 4. DeepCAST-sFDR controls false positives at levels comparable to GWAS.** The heatmap shows the change in false positive rates ($\Delta$ percentage points) relative to GWAS across cohort sizes. While DeepCAST-sFDR yields increases in some settings (red), it also reduces false positives in others (blue). Overall, DeepCAST-sFDR maintains reliability similar to standard GWAS while enabling increased sensitivity.

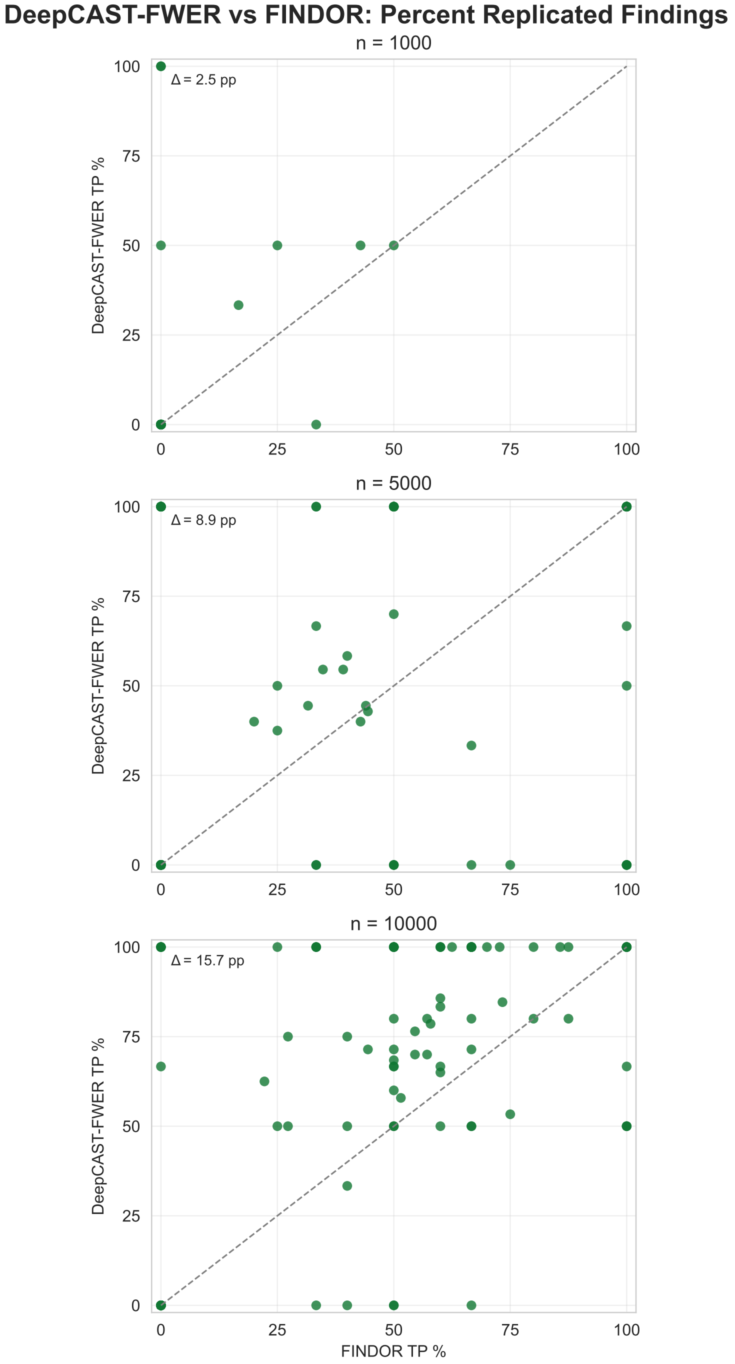


**Fig. 5. DeepCAST-FWER improves replication relative to FINDOR across subsampled cohort sizes.** Scatterplots compare the percentage of lead SNPs that replicate in the full-cohort GWAS for DeepCAST-FWER versus FINDOR at cohort sizes of n = 1000, n = 5000, and n = 10000. Each point represents a replicate of the subsampled phenotype cohort. The diagonal denotes equal performance between methods. Across all sample sizes, most points lie above the diagonal, indicating higher replicated fractions under DeepCAST-FWER. Mean improvements increase with cohort size, from +2.5 percentage points at n = 1000 to +15.7 percentage points at n = 10000.

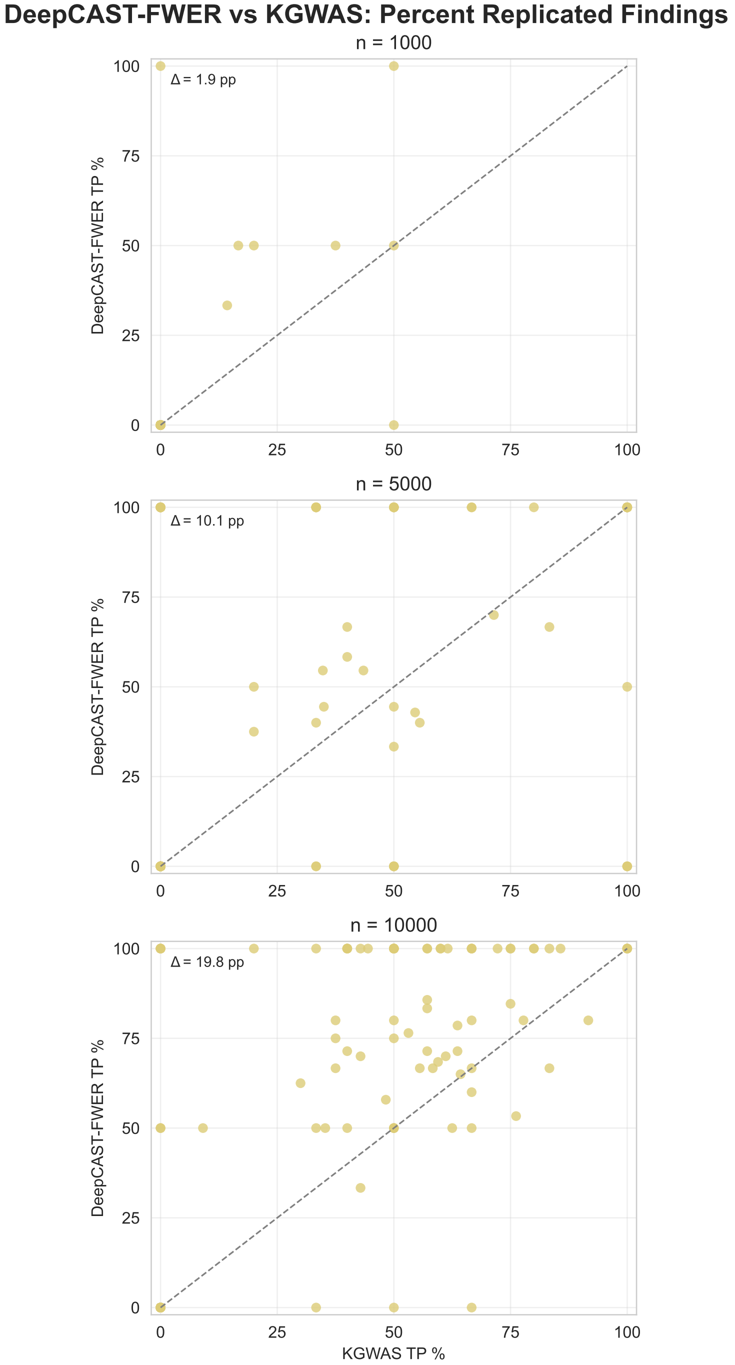


**Fig. 6. DeepCAST-FWER yields higher replicated true positive rates than KGWAS across increasing sample sizes.** Scatterplots show the proportion of lead SNPs replicated in the full-cohort GWAS for DeepCAST-FWER versus KGWAS at n = 1000, n = 5000, and n = 10000. Each point represents a replicate of the subsampled phenotype cohort. Points above the diagonal indicate phenotypes where DeepCAST-FWER produces more reliably replicating associations. Improvements grow with sample size, from +1.9 percentage points at n = 1000 to nearly +20 percentage points at n = 10000, reflecting stronger robustness of DeepCAST-FWER.
